# The Dystrophin-Dystroglycan complex ensures cytokinesis efficiency in *Drosophila* epithelia

**DOI:** 10.1101/2024.03.14.585005

**Authors:** Margarida Gonçalves, Catarina Lopes, Hervé Alégot, Mariana Osswald, Floris Bosveld, Carolina Ramos, Graziella Richard, Yohanns Bellaiche, Vincent Mirouse, Eurico Morais-de-Sá

## Abstract

Cytokinesis physically separates daughter cells at the end of cell division. This step is particularly challenging for epithelial cells, which are connected to their neighbors and to the extracellular matrix by transmembrane protein complexes. To systematically evaluate the impact of the cell adhesion machinery on epithelial cytokinesis efficiency, we performed an RNAi-based modifier screen in the *Drosophila* follicular epithelium. Strikingly, this unveiled adhesion molecules and transmembrane receptors that facilitate cytokinesis completion. Among these is Dystroglycan, which connects the extracellular matrix to the cytoskeleton via Dystrophin. Live imaging revealed that Dystrophin and Dystroglycan become enriched in the ingressing membrane, below the cytokinetic ring, during and after ring constriction. Using multiple alleles, including Dystrophin isoform-specific mutants, we show that Dystrophin/Dystroglycan localization is linked with unanticipated roles in regulating cytokinetic ring contraction and in preventing membrane regression during the abscission period. Altogether, we provide evidence that, rather than opposing cytokinesis completion, the machinery involved in cell-cell and cell-matrix interactions has also evolved functions to ensure cytokinesis efficiency in epithelial tissues.

## INTRODUCTION

Cytokinesis, the physical separation of daughter cells, relies on a dramatic remodeling of the cytoskeleton. It begins with the assembly of an actomyosin contractile ring at the cell equator, which drives membrane ingression, and is completed by the abscission of a small intercellular bridge, or midbody ring, formed at the end of ring constriction. Cytokinesis has fascinated researchers over the past fifty years, leading to the identification of a set of structural components and regulators that are conserved between amoebae, fungi and animals (Pollard & O’Shaughnessy, 2019). These include formins and non-muscle Myosin II, which drive F-actin polymerisation and actomyosin contraction, respectively, and anillin and septins, which tether the ring to the plasma membrane. However, the mechanisms of cytokinetic ring contraction must be adapted to the cell intrinsic and extrinsic specificities of different animal cell types (Cabernard *et al*, 2010; Davies *et al*, 2018; Jordan *et al*, 2016; Ozugergin & Piekny, 2022; Paim & FitzHarris, 2022). A particularly interesting case is provided by epithelia, where extrinsic mechanical forces generated by cell-cell and cell-extracellular matrix (ECM) interactions poses unique challenges during cytokinesis (Herszterg *et al*, 2014; Osswald & Morais-de-Sa, 2019). Cytokinesis failure leads to whole genome duplication and centrosome amplification, which are key contributors to genetic instability and often associated with epithelial oncogenesis (Lens & Medema, 2019; Levine *et al*, 2017; Vittoria *et al*, 2023).

During cytokinesis, epithelial cells remodel cell-cell junctions while maintaining tissue cohesion and the ability to act as a physical and chemical barrier (Daniel *et al*, 2018; Higashi *et al*, 2016; McKinley *et al*, 2018; Wang *et al*, 2018). In line with the prominent role of E-cadherin-based adherens junctions (AJ) in the mechanical coordination between epithelial cells (Lecuit & Yap, 2015), studies of epithelial cytokinesis focused on the understanding of AJ remodeling, the coordination between cytokinesis and *de novo* junction formation, and the role of AJ in the direction of ring closure (di Pietro *et al*, 2023; Firmino *et al*, 2016; Founounou *et al*, 2013; Guillot & Lecuit, 2013; Herszterg *et al*, 2013; Morais-de-Sa & Sunkel, 2013; Pinheiro *et al*, 2017). These aspects of epithelial cytokinesis play a role in the topology and morphogenesis of animal tissues. However, few studies have yet examined the impact of the unique epithelial context on the efficiency of cytokinesis completion. Work in the *Drosophila* pupal notum epithelium has shown that the presence of AJ promotes cytokinesis failure in a background sensitized by loss of septins (Founounou *et al*., 2013). In line with this, increased tension at AJ due to tight junction defects or by excessive neighbor cell contractility led to cytokinesis failure in the *Xenopus* epithelia (Hatte *et al*, 2018; Landino *et al*, 2023). Cell-matrix interactions could also interfere with epithelial cytokinesis and, accordingly, focal adhesions connect the cytokinetic ring to the ECM in the zebrafish epicardium, and promote cytokinesis failure upon adhesion reinforcement (Uroz *et al*, 2019). Thus, while cell adhesion molecules are not inherently essential for cytokinesis completion, their potential to negatively interfere with cytokinesis efficiency could threaten organismal homeostasis.

The follicular epithelium of the *Drosophila* ovary combines genetic tractability with the power to image epithelial cytokinesis in a proliferative adult organ. We designed an RNAi-based *in vivo* genetic modifier screen to test the impact of the major regulators of cell-cell and cell-matrix interactions on cytokinesis efficiency. This uncovered unexpected cytokinetic functions for the Dystrophin-Dystroglycan complex, a transmembrane complex that links the ECM to the cytoskeleton and has important implications in the pathogenesis of neuromuscular dystrophies and cancer (Jones *et al*, 2021; Mirouse, 2023; Nowak & Davies, 2004).

## RESULTS AND DISCUSSION

### *Drosophila* RNAi modifier screen to test the impact of cell-cell and cell-matrix interactions on epithelial cytokinesis efficiency

We performed an RNAi modifier screen in the *Drosophila* follicular epithelium using a modifiable cytokinesis-sensitized background produced by RNAi against Anillin, a component of the cytokinetic ring that tethers the contractile ring and, subsequently, the midbody ring to the plasma membrane (Kechad *et al*, 2012; Zhang & Maddox, 2010). We aimed to ensure temporally-controlled and tissue-specific expression of Anillin RNAi to produce a modifiable phenotype with mild frequency of multinucleated cells. We therefore combined UAS-driven Anillin RNAi induced by a follicular epithelium specific *tj-GAL4* driver (Olivieri *et al*, 2010), with Gal80^ts^ (McGuire *et al*, 2003), a temperature-sensitive repressor of the GAL4 transcription factor, for RNAi induction by incubation at restrictive temperature 3 days prior to tissue dissection (Fig 1A). We used a fluorescent marker of the plasma membrane (Myr:GFP) together with DAPI staining for custom-made automated image quantification of the multinucleation ratio in stage 10 egg chambers (Fig S1), which was used as a proxy for cytokinesis failure during earlier proliferative stages. Under these experimental conditions, Anillin RNAi led to a mild multinucleation level, which allowed us to monitor rescue or enhancement of cytokinesis defects.

**Figure 1.**
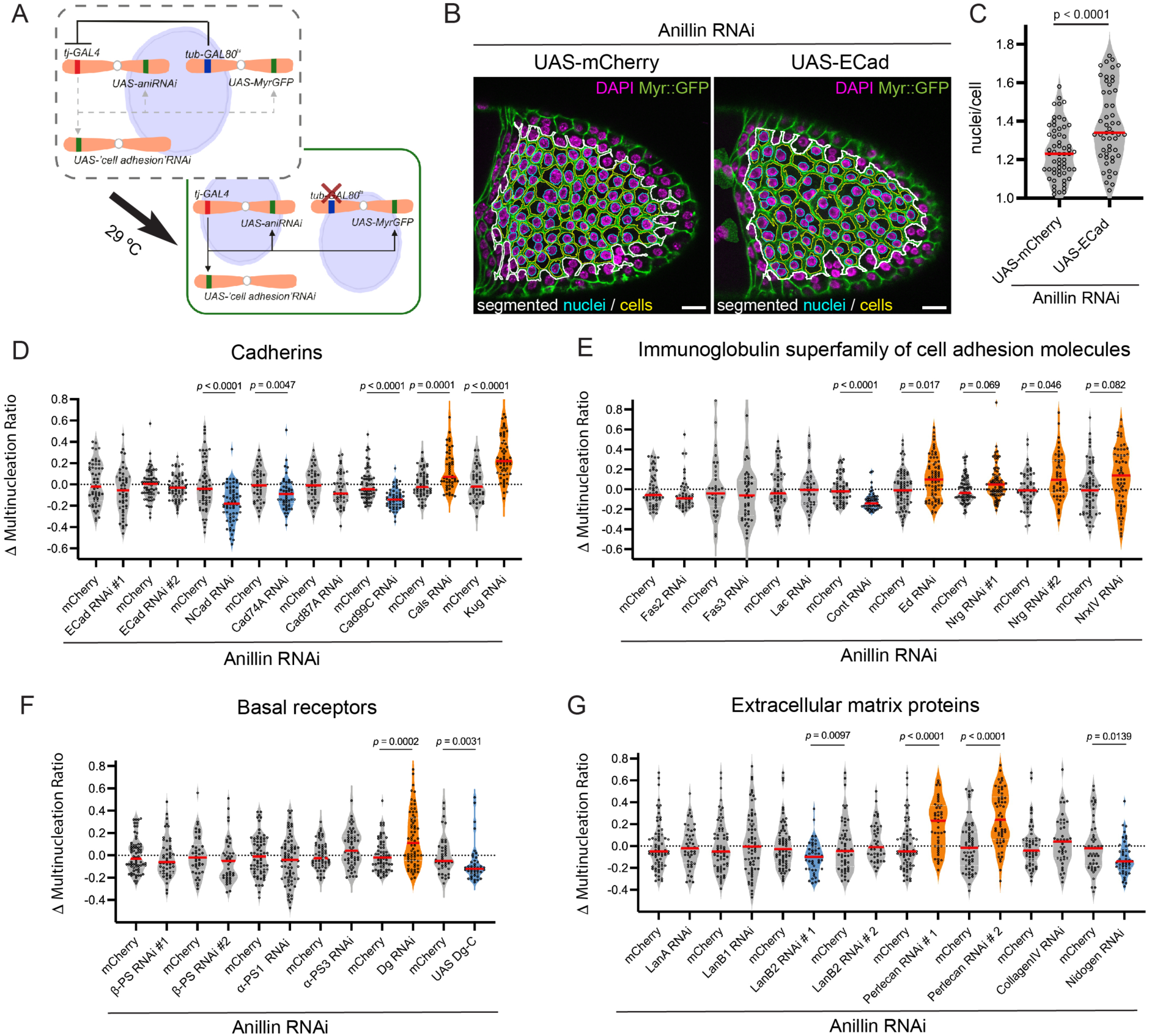
Impact of cell adhesion molecules on epithelial cytokinesis efficiency. **A)** Strategy for the RNAi modifier screen in the *Drosophila* follicular epithelium. Gal80^ts^, a temperature-sensitive repressor of the Gal4 transcription factor, inhibits UAS-RNAi expression at a restrictive temperature. Inactivation of Gal80^ts^ by incubation at 29°C induces protein depletion and expression of a genetically encoded membrane marker (Myr:GFP) in the follicular epithelium. **B)** Automated segmentation of cells (yellow) and nuclei (cyan) in the central area (white line) of egg chambers expressing Anillin RNAi simultaneously with a control transgene (UAS-mCherry) or overexpressing E-cadherin (UAS-ECad). Cell membranes are marked with Myr:GFP and DAPI staining labels the nuclei (magenta). **C)** Ratio of nuclei/cell in egg chambers expressing E-cadherin (UAS-ECad, n = 52 stage 10 egg chambers) and UAS-mCherry (n = 59) in a cytokinesis-sensitized background (Anillin RNAi). Each dot in the violin plot represents the multinucleation ratio of an egg chamber. Median is indicated. *p*-value calculated by non-parametric unpaired Mann-Whitney test. **D-G)** Modification of the Anillin RNAi multinucleated cell phenotype by co-depletion of proteins from the Cadherin (D) and Immunoglobulin superfamily (E), basal receptors (F) and extracellular matrix components (G). Enhancers (orange) and suppressors (blue) are highlighted. For each UAS-driven RNAi, co-expression of an inert UAS-mCherry was used as a control. Each dot in the violin plot represents the Δ Multinucleation Ratio of an analyzed egg chamber in comparison to the mean of the respective control (Δ Multinucleation Ratio = (nuclei/cell)_RNAi_ - mean (nuclei/cell)_mCherry_). Median is indicated. *p*-value calculated by non-parametric unpaired Mann-Whitney test.

To validate the strategy to identify modifiers of cytokinesis efficiency, we tested the impact of reinforcing apical cell adhesion. Overexpression of E-Cadherin (ECad) led to a significant increase of multinucleated cells (Fig 1B and C). This is consistent with previous findings suggesting that AJ challenge cytokinesis completion in *Drosophila* tissues (Founounou *et al*., 2013). We then used UAS-driven RNAi lines to test the impact of the transmembrane proteins of the cadherin superfamily expressed in the follicular epithelium (based on the transcriptional profile of modENCODE (Brown *et al*, 2014)). Multinucleation produced by each RNAi line was compared to the reference value of Anillin depletion on its own, determined for each independent experiment and produced by co-expression of UAS-Anillin RNAi with UAS-mCherry (Δ multinucleation ratio = (nuclei/cell)_RNAi_ - mean (nuclei/cell)_mCherry_). Intriguingly, we observed no effect on cell multinucleation by depletion of E-Cadherin (Fig 1D), whereas depleting N-cadherin (NCad) reduced the number of multinucleated cells. This suggests that, whereas N-Cadherin depletion effectively reduces apical cell adhesion in follicle cells, E-Cadherin depletion does not, which is most likely explained by the fact that NCad and ECad are co-expressed and that N-Cadherin can compensate for E-cadherin loss during the proliferative stages of oogenesis ((Tanentzapf *et al*, 2000) and Fig S2A). Moreover, we found that depletion of two other apical cell adhesion molecules, Cadherin 74A (Cad74A) and Cadherin 99C (Cad99C), also reduced the multinucleation ratio (Fig 1D). In contrast, depletion of the non-canonical cadherin Fat2, *kugelei* (*kug*/*fat2*), which is enriched at the basal side of epithelial cells (Viktorinova *et al*, 2009), or of Calsyntenin (Cals), the localization of which remains to be studied, increased the multinucleation ratio (Fig 1D). Altogether, this suggested that cadherin-based adhesion can oppose cytokinesis completion in the follicular epithelium only when exerted at the apical level.

A second large family of polarized cell adhesion molecules belongs to the immunoglobulin superfamily (Finegan & Bergstralh, 2020). This includes Echinoid (Ed), the *Drosophila* functional orthologue of mammalian Nectin, which forms a second adhesion complex at AJ, as well as Neuroglian (Nrg), Fasciclin 2 (Fas2), Fasciclin 3 (Fas3), Lachesin (Lac) and Contactin (Cont), which are distributed along the lateral membrane of the proliferative follicular epithelium, prior to septate junction maturation. Although we did not observe any effect of depleting Fas2, Fas3 and Lac, we found that Cont RNAi suppressed the Anillin RNAi phenotype, whereas Nrg and Ed enhanced the multinucleation ratio (Fig 1E). Nrg can form heterotypic interactions with Ed, and can also bind the single-pass transmembrane protein Neurexin IV (NrxIV) (Banerjee *et al*, 2006; Islam *et al*, 2003). NrxIV RNAi also increased multinucleation, suggesting that Nrg-NrxIV and Nrg-Ed complexes formed in the lateral membrane could promote epithelial cytokinesis efficiency.

Integrins are the central physical linkers between the cytoskeleton and the extracellular matrix that work as heterodimeric receptors formed by an α and a β subunits (Kanchanawong & Calderwood, 2023). However, knockdown of the integrin gene encoding Myospheroid (*mys/*βPS), the only β subunit expressed in follicle cells, or of the α subunits Scab (*scb*/αPS3) and Multiple edematous wings (*mew*/αPS1) did not produce any effect on the multinucleation ratio (Fig 1F). To corroborate this conclusion, we confirmed the protein depletion of βPS by immunofluorescence and we validated the functional impact of RNAi-mediated depletion of integrins (Fig S2B and E), showing that egg chamber elongation is affected as previously reported (Qin *et al*, 2017). We then tested the impact of the non-integrin ECM receptor Dystroglycan (*Dg*), which also plays a role in *Drosophila* egg morphogenesis (Cerqueira Campos *et al*, 2020; Schneider *et al*, 2006). Knockdown Dg increased multinucleation, suggesting that it plays a positive role in epithelial cytokinesis.

The mechanical properties of the matrix and its ability to regulate signaling from focal adhesions modulates cytokinesis efficiency in cell culture models (Rabie *et al*, 2021; Sambandamoorthy *et al*, 2015). We thereby postulated that ECM composition could impact cytokinesis efficiency and screened for the impact of the main ECM components expressed in the basement membranes of follicle cells, namely Collagen IV (*viking*/ Col4α2), Laminins, Perlecan and Nidogen (Diaz-Torres *et al*, 2021). Depletion of Collagen IV, the major regulator of basement membrane stiffness in the follicular epithelium (Crest *et al*, 2017; Topfer *et al*, 2022), caused defects on egg chamber elongation (Fig S2E), but did not modify the multinucleation ratio of Anillin RNAi (Fig 1G and S2C). Depletion of Nidogen and Laminin B2 (LanB2) produced a significant reduction of multinucleation, but RNAi for Laminin A (LanA) did not modify the multinucleation ratio, despite of an efficient reduction of protein levels (Fig S2D). In contrast, Perlecan (*trol*) depletion with distinct RNAi lines led to a dramatic increase in multinucleation, suggesting this protein promotes cytokinesis robustness (Fig 1G).

Altogether, the genetic modifier screen indicates that whereas AJ mainly challenges epithelial cytokinesis efficiency in follicle cells, there are a number of transmembrane proteins involved in cell-cell and cell-matrix interactions that promote efficient cytokinesis (overview of screen data in Table S1). It is worth noting that depletion of Ed, Nrg, NrxIV, Dg, Kug or Perlecan did not produce multinucleated cells on their own (Fig S2F), which suggests that these proteins contribute to cytokinesis efficiency in a sensitized context, but are dispensable for cytokinesis completion.

### The Dystrophin-Associated Protein Complex promotes cytokinesis efficiency

The importance of regulating cell-substrate interactions has been well studied by cell cultures studies (Taneja *et al*, 2019), but remains unexplored in epithelial tissues. We therefore addressed the putative role for the basal ECM receptor Dg in cytokinesis efficiency. To further test Dg function, we analyzed the impact of overexpressing Dg. Dg overexpression reduced the cytokinesis defects caused by Anillin RNAi (Fig 1F), providing additional evidence for the positive role of Dg on epithelial cytokinesis efficiency. Dg is a transmembrane heterodimeric protein that is part of the Dystrophin-associated protein complex (DAPC). It links the ECM to the intracellular cytoskeleton by interacting with Dystrophin, an actin-binding protein that binds the cytoplasmic tail of Dg (Fig 2A). Intriguingly, the human orthologues of Dystroglycan and Dystrophin (Dp71) are enriched in the cleavage furrow and the midbody in *in vitro* cell culture models (Higginson *et al*, 2008; Villarreal-Silva *et al*, 2011), yet their role in cytokinesis remains unknown.

**Figure 2.**
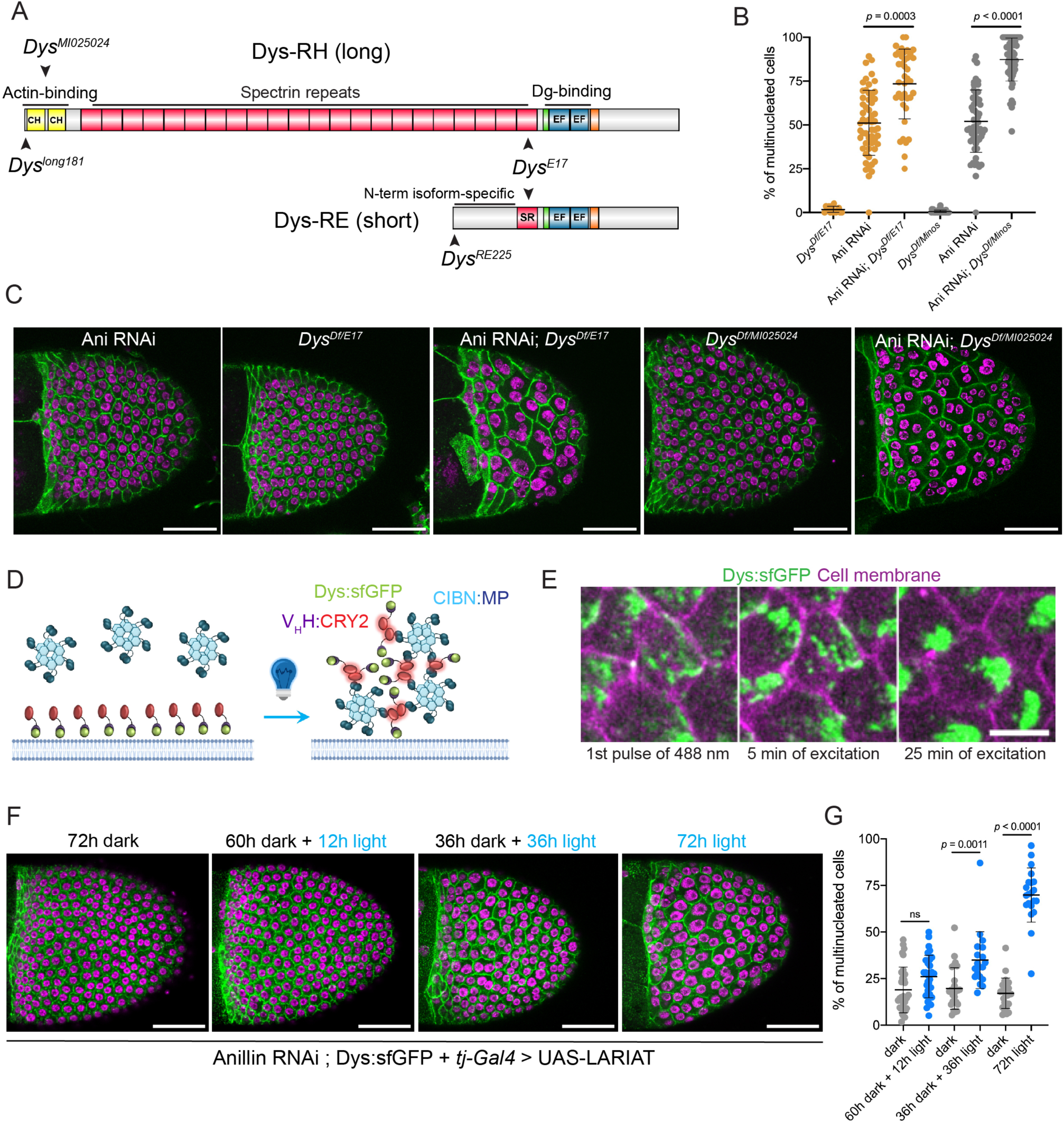
Dystrophin promotes cytokinesis efficiency. **A)** Schematic representation of Dystrophin isoforms expressed in the *Drosophila* follicular epithelium. Dys-RH (long isoform) encodes a large protein containing an N-term actin-binding domain, a rod domain comprised of several spectrin repeats and a C-term Dystroglycan-binding domain. Dys-RE (short isoform) lacks the N-term actin-binding domain and the majority of the rod domain. The position of the truncating mutations for *Dys^E17^,* which disrupts the Dg-binding domain of both isoforms, *Dys^long181^,* which specifically disrupts the long isoform, and *Dys^RE225^*, which specifically disrupts the short isoform, is shown. *Dys^MI025024^*is a *minos* insertion that disrupts the actin-binding domain of the long isoform. **B)** Frequency of multinucleated cells in stage 10 egg chambers from Anillin RNAi, *Dys^E17/Df^* and *Dys^MI025024/Df^* mutants in unperturbed or sensitized (Anillin RNAi) cytokinetic background. Graph shows mean ± SD. Each dot represents the multinucleation ratio of an egg chamber. *p*-value calculated by non-parametric unpaired Mann-Whitney test. **C)** Representative surface images of the egg chambers quantified in B, expressing ECad:GFP to contour the cell membranes (green) and DAPI staining to label nuclei (magenta). Scale bars: 50 µm. **D)** Schematic representation of light-induced clustering of Dys:sfGFP. The system includes a GFP nanobody (V_H_H) fused with CRY2 to recruit endogenously tagged Dys:sfGFP and CIBN fused with a multimerization domain (MP). Blue light exposure triggers cluster formation through light-induced homodimerization of CRY2 and heterodimerization of CRY2 with CIBN. **E)** Time-lapse images of endogenously tagged Dys:sfGFP (green) egg chambers co-expressing LARIAT (basal view) and stained with the membrane marker (cell mask, magenta). Imaging with the 488 nm laser triggers fast reallocation of Dys:sfGFP to large single clusters within each cell. Scale bar: 5 µm. **F)** Representative images of endogenously tagged Dys:sfGFP stage 10 egg chambers co-expressing LARIAT and Anillin RNAi from flies exposed to the indicated periods of light. A control group was kept in the dark for 72h in parallel with each experimental condition. Egg chambers were stained for Armadillo to contour the cell membranes (green) and DAPI to label the nuclei (magenta). Scale bars: 50 µm. **G)** Frequency of multinucleated cells from the experimental conditions represented in F. Graph shows mean ± SD. Each dot represents the multinucleation ratio of an egg chamber. *p*-value calculated by non-parametric unpaired Mann-Whitney test.

To investigate if Dys also modulates epithelial cytokinesis, we assessed the genetic interaction between Anillin RNAi and *Dys* mutant alleles. We used a genomic deletion (*Df (3R)Exel6184*, named *Dys^Df^*) that spans the entire Dys locus, along with *Dys^E17^* (nonsense mutation resulting in truncated protein that disrupts the C-term Dg-binding domain) or with *Dys^MI025024^* (*minos* insertion disrupting the exon encoding the actin-binding domain of the long Dys isoforms) (Fig 2A). Neither *Dys^E17^*^/*Df*^ nor *Dys^MI025024^*^/*Df*^ egg chambers exhibited a significant number of multinucleated cells, suggesting that Dys is dispensable for cytokinesis completion (Fig 2B and C). However, disruption of *Dys* function using both mutant alleles in the background of Anillin RNAi led to a dramatic increase in the frequency of multinucleated cells when compared to Anillin RNAi alone (Fig 2B and C).

Multinucleation could be generated independently of proliferation by cell fusion events (Bailey *et al*, 2021). To further validate Dys role in cytokinesis efficiency, we used an optogenetic approach (light-activated reversible inhibition by assembled trap (LARIAT)) that enables temporally-controlled disruption of protein function by light-induced clustering of endogenously GFP-tagged proteins (Osswald *et al*, 2022; Qin *et al*., 2017). To target Dys, we used flies with endogenously sfGFP:tagged Dys, which labels all Dys isoforms (Dennis *et al*, 2023), and co-expressed GAL4-driven UAS-LARIAT (UAS-V_H_H:CRY2-P2A-CIBN:MP) in the follicular epithelium (Fig 2D). We first confirmed that blue light exposure of egg chambers cultured *ex vivo* quickly induces Dys:sfGFP clustering in the basal cortex of follicle cells (Fig 2E). We then tested if exposure of living flies during specific periods of light necessary to cluster Dys:sfGFP in proliferative and/or non-proliferative stages would modify the frequency of multinucleated identified in stage 10 upon Anillin RNAi. Exposing flies during 12h to blue light, which only allows progression from non-proliferative stage 7 during light exposure, did not modify the frequency of multinucleated cells (Fig 2F and G). Increased periods of light exposure (36h and 72h) to ensure Dys perturbation during proliferation (stages 1-6 of oogenesis) led to a dramatic increase in the frequency of multinucleated cells. These results suggest that the increased multinucleation of follicle cells quantified in stage 10 egg chambers is associated with cytokinesis failure during the proliferative stages. Together, we conclude that the DAPC plays a protective role against cytokinesis failure.

### Dynamic distribution of the DAPC during cytokinesis in *Drosophila* epithelia

To investigate how the DAPC functions during epithelial cytokinesis, we examined its distribution by time-lapse movies of follicle cell division. Using an UAS-driven Dg:GFP in egg chambers co-expressing Anillin:RFP, to label the cytokinetic ring (Fig 3A), we observed that Dg:GFP strongly accumulated at the ingressing membrane below the cytokinetic ring. After ring closure, Dg:GFP accumulated near the midbody. Overexpressed Dg:GFP was not restricted to the basal side in interphase cells, contrary to previous observations by antibody staining (Schneider *et al*., 2006). To rule out protein overexpression effects on the localization of the DAPC during cell division, we examined the mitotic redistribution of endogenously tagged Dg:GFP and Dys:sfGFP lines (Fig 3B and C, Movie S1-S3). Endogenously tagged Dys:GFP and Dg:GFP were largely restricted to the basal domain during interphase, and also become enriched at the plasma membrane during cell rounding and at the ingressing membrane during ring constriction. After ring constriction, both proteins accumulated in the new interface formed between daughter cells, close to the midbody ring (Fig 3B-D). To evaluate if the mitotic redistribution of Dys-Dg could be generalized to other proliferative tissues, we imaged the *Drosophila* thorax epithelium (notum) of the fly pupa. Dg and Dys also became strongly enriched at the ingressing membrane just below the contractile ring in this tissue, and persisted along the new membrane below the midbody ring post-constriction (Fig S3). Together, this shows that the Dg-Dys complex exhibits dynamic membrane redistribution during cytokinesis in different tissues, in line with a role during epithelial cytokinesis.

**Figure 3.**
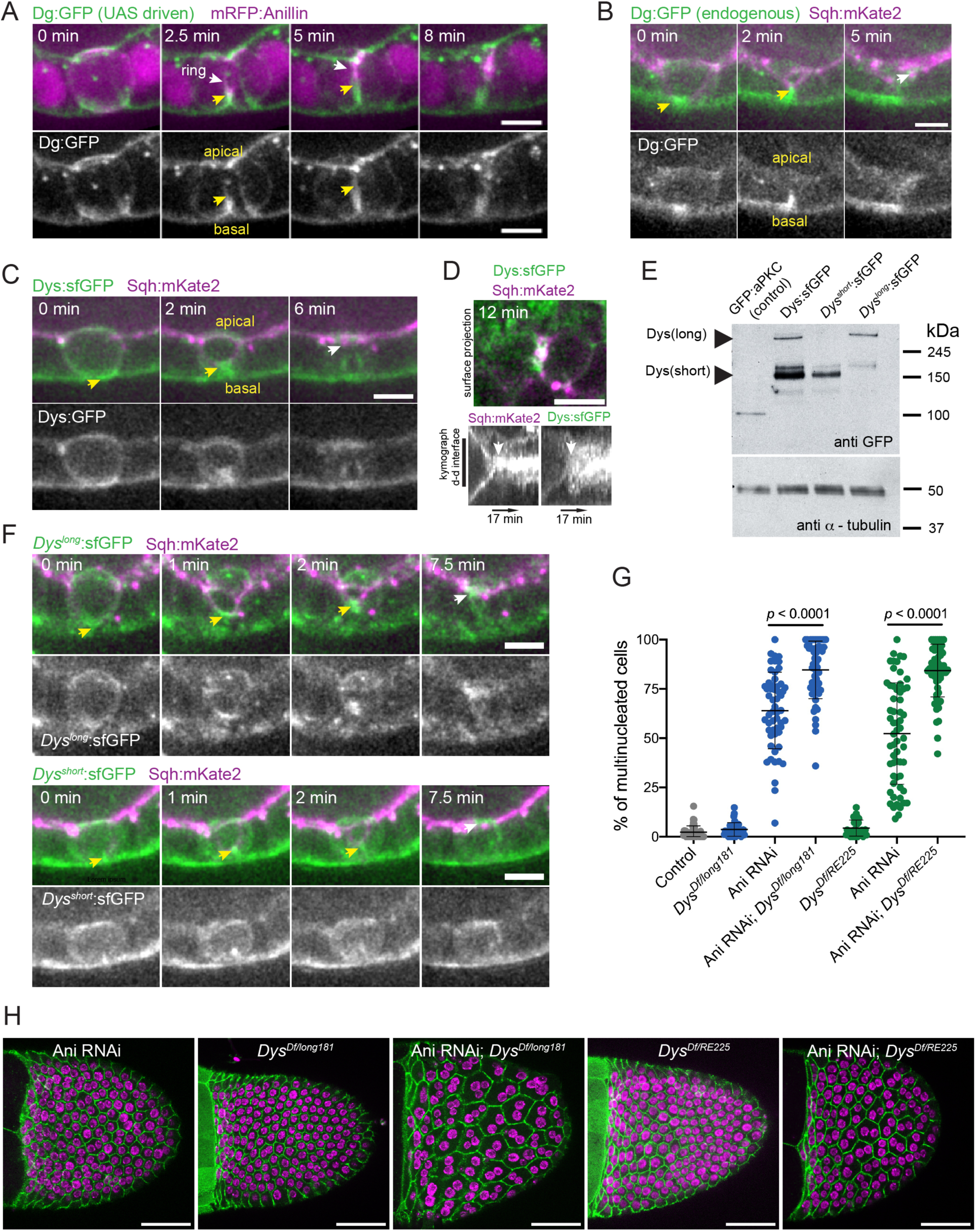
Dynamic redistribution of the DAPC during epithelial cytokinesis. **A)** Time-lapse images of follicle cells expressing UAS-driven Dg:GFP and Anillin:mRFP. Projection along the apico-basal axis of the epithelia shows Dg:GFP enrichment in the ingressing cleavage furrow (yellow arrows). Scale bars: 5 µm. **B,C)** Accumulation of endogenously tagged Dg:GFP (B) and Dys:sfGFP (C) at the basal part of the ingressing membrane since the beginning of ring (labelled with Sqh:mKate2) constriction (yellow arrows). After ring closure, Dg:GFP (B) and Dys:sfGFP (C) show an enrichment close to the midbody (white arrows). Scale bars: 5 µm. **D)** Follicle cells surface projection from a time-lapse movie showing the localization of Dys:sfGFP (green) at the new daughter cell interface. A related kymograph for a line along the daughter-daughter (d-d) interface shows that, after ring closure followed with Sqh:mKate2 distribution (white arrow), there is an enrichment of Dys:sfGFP at the new d-d interface. Scale bars: 5 µm. **E)** Western blots of protein extracts from GFP:aPKC, Dys:sfGFP; *Dys^short^*:sfGFP and *Dys^long^*:sfGFP flies probed with anti-GFP or α-tubulin as loading control. GFP:aPKC was used as control of GFP detection. **F)** Accumulation of endogenously tagged *Dys^long^*:sfGFP (top panel) and *Dys^short^*:sfGFP (bottom panel) at the basal part of the ingressing membrane since the beginning of ring (labelled with Sqh:mKate2) constriction (yellow arrows). After ring closure, *Dys^long^*:sfGFP and *Dys^short^*:sfGFP show an enrichment close to the apically positioned midbody (white arrows). **G)** Frequency of multinucleated cells in stage 10 egg chambers from control, *Dys^long181/Df^*and *Dys^RE225/Df^* mutants in unperturbed or sensitized (Ani RNAi) cytokinetic background. Graph shows mean ± SD. Each dot represents the multinucleation ratio of an egg chamber. *p*-value calculated by non-parametric unpaired Mann-Whitney test. **H)** Representative surface images of the egg chambers quantified in (G), expressing ECad:GFP to contour the cell membrane (green) and DAPI staining to label nuclei (magenta). Scale bars: 50 µm. A,B,C,D,F - Time is indicated relative to the beginning of constriction.

Dys encodes several isoforms in both humans and flies. All of them bear a conserved C-terminal region containing the Dystroglycan binding domain, but the shorter isoforms lack the known actin-binding domains (N-term actinin like domain and a second actin-binding domain within the rod domain). Translating ribosome affinity purification experiments show that only one long (Dys-RH) and one short isoform (Dys-RE) are expressed in the follicular epithelium (Vachias *et al*, 2023). To investigate whether the redistribution and function of Dys during cytokinesis relies on its ability to directly bind the actin cytoskeleton, we generated isoform-specific mutant alleles in an untagged version of the *Dys* genomic locus *(Dys^long181^,* i.e. mutating all the long isoforms including RH, and *Dys^RE225^* mutating RE) or in a sfGFP-tagged one (*Dys:*sfGFP*^long181^* and *Dys:*sfGFP*^RE225^*), by CRISPR/Cas9-mediated introduction of indel mutations. As follicle cells normally express only a short and a long isoform, we simplified the naming of *Dys:*sfGFP*^long181^* and *Dys:*sfGFP*^RE225^* genotypes by indicating the tagged isoforms *Dys^short^*:sfGFP and *Dys^long^*:sfGFP, respectively. Accordingly, western blot of ovary extracts with anti-GFP antibodies confirm that the long isoform is not produced in *Dys^short^*:sfGFP and that the short isoform is absent from *Dys^long^*:sfGFP (Fig 3E). Thus, these lines were used to evaluate the relative contributions of short and long isoforms on Dys distribution. Both Dys isoforms showed strong basal enrichment in the proliferative follicular epithelium. However, *Dys^short^*:sfGFP exhibited a more uniform distribution in the basal cortex that contrasts with a highly planar polarized distribution of *Dys^long^*:sfGFP (Fig S4). More importantly, both isoforms showed some local enrichment below the ingressing furrow during constriction, and at the new interface post-constriction, suggesting that both contribute to Dys role in epithelial cytokinesis (Fig 3F and Movie S4). Accordingly, disruption of *Dys* function using either one of the isoform-specific mutant alleles led to a dramatic increase of multinucleated cells in tissues depleted of Anillin by RNAi (Fig 3G and H). Thus, both Dys isoforms, regardless of the presence of actin-binding domains, localize at the ingression furrow and contribute to epithelial cytokinesis efficiency.

### Multiple roles of the DAPC complex on epithelial cytokinesis efficiency

Anillin is required for the maturation of the midbody ring that stabilizes the intercellular bridges during abscission, and accordingly, Anillin RNAi in *Drosophila* S2 cells prevalently leads to reopening of the furrow a significant period of time after ring constriction (Kechad *et al*., 2012). To further understand the genetic interaction between Anillin RNAi and *Dys* in epithelial cytokinesis, we monitored follicle cell cytokinesis by live imaging (Fig 4A). We could barely detect cytokinesis failure when cells were depleted of Anillin on its own, while perturbation of *Dys* function increased the frequency of cells failing cytokinesis (Fig 4C). Although the vast majority of cells reached a semi-stable state after complete ring closure, the new interface formed between the daughter cells regressed as long as 30 minutes post ring constriction (Fig 4B and C and Movie S5). Thus, we provide evidence that the DAPC contributes to the efficiency of the last stage of cytokinesis.

**Figure 4.**
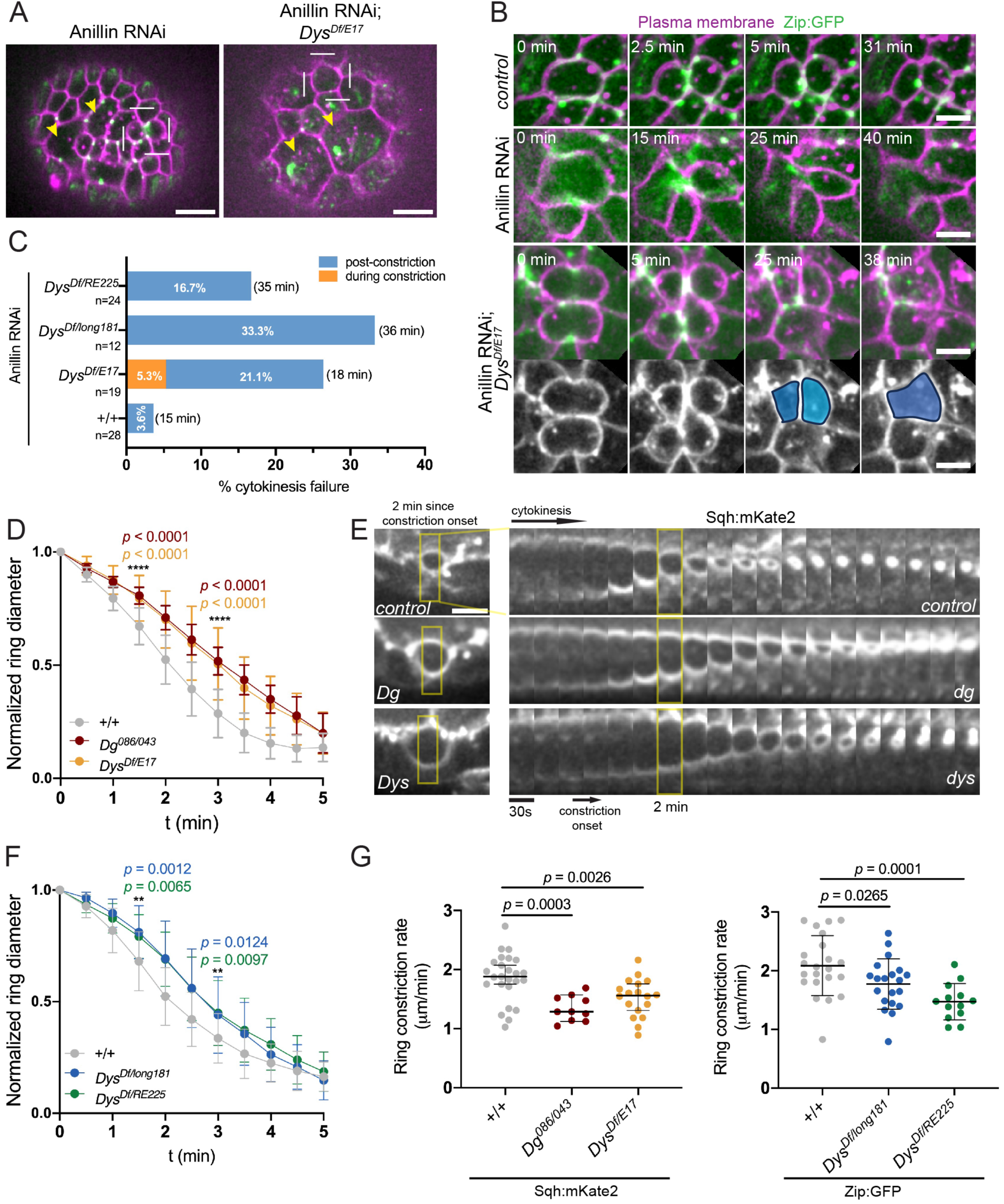
Multiple roles of the Dg-Dys complex in epithelial cytokinesis efficiency. **A)** Surface view of Anillin RNAi and *Dys*, Anillin RNAi egg chambers. Both conditions express Zip:GFP (green) and were membrane stained (CellMask, magenta). Note the presence of large cells produced by cytokinesis failure (yellow arrows). Quantification of the cytokinesis defects by live imaging (B, C) was restricted to cells without previous cytokinesis failure (white box). Scale bars: 10 µm. **B)** Time-lapse images (surface view) of Zip:GFP (green) egg chambers stained with a membrane marker (magenta), captured from control, Anillin RNAi and *Dys*, Anillin RNAi. Disruption of *Dys* function in follicle cells depleted of Anillin leads to cytokinesis failure due to membrane regression after closure of the contractile ring. Scale bars: 5 µm. **C)** Frequency of cytokinesis failure in egg chambers only depleted of Anillin (+/+) or with concomitant disruption of *Dys* function using the indicated allelic combinations. For the post-constriction failure, the time elapsed (average) since the end of ring constriction until membrane regression is shown. **D)** Contractile ring diameter (mean ± SD, normalized to initial value) during follicle cell cytokinesis in control (n = 25), *Dg^086/043^* (n = 10) and *Dys^E17/Df^* (n = 18) mutant egg chambers. The indicated *p*-values correspond to the difference in ring diameter between *Dg^086/043^* (red) or *Dys^E17/Df^* (yellow) and control egg chambers, at *t*=1.5 and *t*=3 min. **E)** Time-lapse of cytokinetic ring constriction viewed along the apical-basal axis in control, *Dg^086/043^* and *Dys^E17/Df^* mutant egg chambers expressing Sqh:mKate2, 2 minutes since constriction onset. Sequential projections of ring constriction for all genotypes are also shown. Scale bars: 5 µm. **F)** Contractile ring diameter (mean ± SD, normalized to initial value) during follicle cell cytokinesis in control (n = 24), *Dys^long181/Df^* (n = 20) and *Dys^RE225/Df^* (n = 13) mutant egg chambers. The indicated *p*-values correspond to the difference in ring diameter between *Dys^long181/Df^* (blue) or *Dys^RE225/Df^* (green) and control egg chambers, at *t*=1.5 and *t*=3 min. **G)** Ring constriction rate during the linear phase of constriction in control (n = 25 / n = 22), *Dg^086/043^* (n = 10), *Dys^E17/Df^* (n = 18), *Dys^long181/Df^* (n = 20) and *Dys^R225/Df^* (n = 13) mutant egg chambers, either expressing Sqh:mKate2 (left) or Zip:GFP (right). Mean ± SD is indicated. *p*-value calculated using non-parametric unpaired Mann-Whitney test.

In line with Dys and Dg spatial redistribution in the furrowing membrane during cytokinesis, we also postulated that the DAPC could regulate early cytokinetic furrowing. We therefore monitored cytokinetic ring constriction using time-lapse movies of *Dg (Dg^043/086^*) and *Dys (Dys^E17/Df^)* mutant egg chambers. Disruption of either *Dg* and *Dys* significantly delayed ring constriction (Fig 4D and E and Movie S6). The delay in ring closure starts during the initiation of furrowing, as there was already a significant difference in ring diameter at the onset of the linear phase of ring constriction (1.5 min). To specifically examine the impact of long and short Dys isoforms, we also imaged mutant egg chambers for each isoform (*Dys^RE225/Df^* or *Dys^long181/Df^*). Both mutants impaired cytokinetic closure, similarly to *Dg* and *Dys* mutants (Fig 4F). In addition, both *Dys* isoforms and *Dg* also contributed to the normal rate of ring constriction measured during the linear phase of ring constriction (Fig 4G). Hence, we conclude that the DAPC ensures normal actomyosin ring constriction in the follicular epithelium.

In conclusion, this study provides new insights into the mechanisms that ensure the efficiency of epithelial cytokinesis, and uncovers new cytokinetic functions for the DAPC in *Drosophila* tissues. Previous *in vivo* studies have shown that cell-cell and cell-matrix adhesion could potentiate cytokinesis failure and delay ring constriction in multiple vertebrate and invertebrate tissues (Founounou *et al*., 2013; Hatte *et al*., 2018; Higashi *et al*., 2016; Uroz *et al*., 2019). The results of our RNAi modifier screen now identify a number of proteins involved in cell-cell and cell-matrix interactions that promote, rather than oppose, cytokinesis efficiency. These include Perlecan, a Dg ligand (Talts *et al*, 1999), and Fat2, which interacts genetically with Dys (Cerqueira Campos *et al*., 2020), as well as cell-cell adhesion molecules from the immunoglobulin superfamily. Building on this screen, we have provided evidence that the DAPC, acting through long and short Dys isoforms, contributes to the efficient daughter cell separation after ring constriction, and also functions during cytokinetic furrowing.

Exactly how the DAPC contributes to each step of cytokinesis remains an important open question. We show that Dystroglycan and Dystrophin become enriched under the contractile ring during furrowing in different epithelial tissues, and so we could postulate that DAPC redistribution contributes to the remodeling of cell-matrix interactions, to facilitate the invagination of the plasma membrane at the basal side. Indeed, the DAPC plays important roles in the dynamic organization of the basement membrane and the underlying cytoskeleton to contribute to different morphogenetic processes (Buisson *et al*, 2014; Cerqueira Campos *et al*., 2020; Villedieu *et al*, 2023). Later on, the enrichment of the DAPC near the apical midbody ring during the final steps of cytokinesis suggests that it may act as a local regulator of the machinery involved in the physical cut or the stabilization of the intercellular bridge during abscission. Interestingly, Dystrophin binds microtubules and pauses their polymerization (Belanto *et al*, 2014; Prins *et al*, 2009), so it could be involved in the disassembly of midbody-associated microtubules. Alternatively, the DAPC may facilitate cytokinesis completion by enhancing cell-matrix adhesion through its ability to link the intracellular cytoskeleton to the ECM. This is consistent with mammalian cell culture studies suggesting that maintaining adhesive contacts with the substrate facilitates cytokinesis completion in cells with a compromised actomyosin ring (Dix *et al*, 2018). However, there is no direct evidence for adhesive properties of the DAPC in epithelia, and the finding that Dys cytokinetic functions require isoforms that lack the known actin-binding domains argues against the importance of a direct link between the DAPC and the actin cytoskeleton. Future work can now focus on these hypotheses to elucidate how the DAPC coordinates the interplay between the intracellular cytoskeleton and the extracellular ECM to promote cytokinesis fidelity, and to address how these functions are integrated in different epithelial tissues.

## MATERIALS AND METHODS

The list of reagents used in this study is found in table S4.

### *Drosophila* lines and maintenance

Fly stocks and genetic crosses were raised in standard fly media (cornmeal/agar/molasses/yeast) at 18°C or 25°C, with 60% humidity and 12h/12h dark light cycle, unless stated otherwise. The details of the fly lines used throughout this study are listed in Table S2. A detailed list of the fly genotypes for each experiment can be found in Table S3. We used *traffic jam-Gal4* to drive the expression of UAS constructs in the *Drosophila* follicular epithelium.

### *In vivo* genetic modifier screen

For the genetic modifier screen, the following fly stock was generated: *tj-Gal4*, UAS-Anillin RNAi/CyO; UAS-Myr:GFP, tub-Gal80^ts^/TM6. The RNAi lines for the analyzed cell adhesion molecules were obtained from *Drosophila* Bloomington Stock Center (DBSC), and are listed in detail in Table S1. To fully suppress premature UAS-driven RNAi expression, we used Gal80^ts^, the temperature-sensitive repressor of Gal4, and kept the crosses at 18°C. 0-4 days after hatching, adult offspring was transferred to 29°C for 3 days, to boost the efficiency of RNAi depletion. After these 3 days, ovaries were dissected and fixed. At least 3 fully independent experiments were performed for each RNAi line used and, for each of these experiments, a minimum of 10 egg chambers (randomly selected from a pool of 6-8 dissected animals) were analyzed.

### Molecular biology and transgenesis

Isoform-specific mutants were generated by CRISPR. gRNAs GGAGGAGCTGAATCTGCAGG and GTGGAGCTGCTCCTCCGACG, targeting long (RA,C,F,G,H,I and K) and RE isoforms, respectively, were cloned in PCFD6 vector and transgenic lines were generated at attP40 landing site. Then, these lines were crossed with a line expressing Cas9 in the germline and with or without the sfGFP KI at *Dys* locus. Indel mutations were isolated in the progeny by sequencing. Alleles selected for this work were *Dys^long181^* (4bp deletion, STOP at amino acid position 181 on RH), *Dys^RE225^* (2bp deletion, STOP at amino acid position 225 on RE), *Dys:*sfGFP*^long181^*(4bp deletion, STOP at amino acid position 181 on RH) and *Dys:*sfGFP*^RE225^* (deletion of 2 bp, STOP at amino acid position 227).

### Optogenetics in the follicular epithelium

For the optogenetic perturbation of Dystrophin, we combined Dys:sfGFP in its endogenous locus with UAS-LARIAT. Crosses were maintained at 18°C and, to avoid optogenetic system activation by visible light, the vials were kept in the dark, inside cardboard boxes and handled in a dark room. For *ex vivo* optogenetic experiments (Fig 2E), ovaries were dissected in the dark using a 593 nm LED light source (SuperBrightLEDs). The CIBN-CRY2 interaction was only triggered upon activation of the 488 nm laser used for GFP-tagged protein imaging.

For *in vivo* optogenetic experiments (Fig 2F and G), adult offspring of the genotypes indicated in Fig 2F and which developed at 18°C was transferred to 25°C for 3 days (note that optogenetic experiments could not be performed at 29°C and, so, as anticipated for milder expression of Anillin RNAi at 25°C, the phenotypes of Anillin RNAi on its own are milder in Fig 2G than in Fig 2B). Flies were continuously exposed to blue light (427 nm LED bulb (SuperBrightLEDs)) for different periods of time – 72h, 36h and 12h. For each replicate, control flies were kept in the dark, and dissected in a dark room. After the 3 days at 25°C, ovaries were dissected and immunostained for Armadillo to label the junctional cortex in order to detect multinucleated cells.

### Fixation and staining of egg chambers

*Drosophila* ovaries were dissected in Schneider’s Insect Medium (Sigma-Aldrich) supplemented with 10% FBS (fetal bovine serum, heat inactivated; Thermo Fisher) and fixed using a 4% paraformaldehyde solution (prepared in PBS with 0.05% Tween 20 (Sigma-Aldrich)) for 20 minutes. After washing three times for 10 minutes with PBT, samples were mounted with Vectashield Mounting Medium with DAPI (Vector Laboratories). Alternatively, for antibody staining, after the post-fixation washes, egg chambers were blocked for 1 hour at room temperature with 10% BSA prepared in PBT and incubated overnight at room temperature with the primary antibody diluted in PBT + 1% BSA. Samples were then washed with PBT + 1% BSA and incubated again for at least two hours at room temperature with the secondary antibody diluted in PBT + 0.1% BSA. After three washing steps with PBT, samples were mounted with Vectashield with DAPI (Vector Laboratories). For F-actin staining, we added Phalloidin-TRITC (Merck, 1:250) to the fixative solution and increased the incubation time to 30 min. The following primary antibodies were used: rat anti-N-cadherin (DSHB DN-Ex #8, 1:50), mouse anti-βPS-integrin (DSHB CF.6G11-s, 1:10) and mouse anti-Armadillo (DSHB N2.7A1, 1:100). Respectively, the following secondary antibodies were used: goat anti-rat Alexa 568 (Invitrogen A11077, 1:300), goat anti-mouse Alexa 488 (Invitrogen A11029, 1:300) and goat anti-mouse Alexa 568 (Invitrogen A11031, 1:300).

### Imaging

Images of fixed *Drosophila* egg chambers were acquired on an inverted laser scanning confocal microscope Leica TCS SP5 II (Leica Microsystems), with HC PL APO CS 20x/0.70 NA water, 40x/1.10 NA water or 63x/1.30 NA glycerine objectives, using the LAS 2.6 software. For live imaging of *Drosophila* egg chambers, individual ovarioles were dissected in *ex vivo* culture medium (Schneider’s medium (Sigma-Aldrich) supplemented with 10% FBS (fetal bovine serum, heat inactivated; Thermo Fisher) and 200 µg/µL insulin (Sigma-Aldrich)) and the enveloping muscle removed. Ovarioles were transferred to new culture medium and imaged on glass bottom dishes (MatTek; No 1.5; P35G-1.5-7-C) with an Andor XD Revolution Spinning Disk Confocal system equipped with two solid state lasers – 488nm and 561nm –, an iXonEM+ DU-897 EMCCD camera and a Yokogawa CSU-22 unit built on an inverted Olympus IX81 microscope with a PLAPON 60x/1.42 NA objective, using iQ software (Andor). When indicated in the figures, to mark the cell membrane, ovarioles were stained with CellMask Orange Plasma membrane Stain (ThermoFisher; diluted 1:10 000 in culture medium) for 15 minutes and washed with *ex vivo* culture medium before imaging. Live imaging was performed at 25°C, with the exception for the experiments where UAS-Anillin RNAi was expressed, which were performed at 29°C. Midsagittal egg chamber cross-sections were used to image the follicular epithelium along the apical-basal axis and z-stacks at the surface of the egg chamber to cross-section the follicular epithelium along the apical-basal axis. Z stacks were collected with serial optical sections separated by 1 µm.

Live imaging of *Drosophila* pupa was conducted as described in (Bosveld *et al*, 2012), during the first round of cell division in the anterior scutum region of the notum epithelium. Pupa imaging was performed using an inverted spinning disk wide homogenizer confocal microscope (CSU-W1, Roper/Zeiss) equipped with a sCMOS camera (Orca Flash4, Hamamatsu) and using a 63x/1.4 NA oil DICII PL APO objective. A 40 slices Ζ-stack was collected with serial optical sections separated by 0.5 µm, and captured every 30 seconds. To obtain apical-basal side views of cytokinesis, a maximum projection was generated from a 2 µm resliced region, centered at the cytokinetic ring.

### Protein extracts and Western blot

We prepared protein extracts from *Drosophila* ovaries (at least 25 flies per genotype) of endogenously-GFP-tagged *Dys*^long^ and *Dys*^short^ mutants, as well as Dys:sfGFP. Protein extracts of endogenously GFP-tagged aPKC were used as a control to detect an unrelated GFP-tagged protein. Dissected ovaries were transferred to lysis buffer (150mM KCl, 75mM HEPES pH 7.5, 1.5 mM EGTA, 1.5mM MgCl2, 15% glycerol, 0.1% NP-40, 1x protease inhibitors cocktail (Roche) and 1x phosphatase inhibitors cocktail 3 (Sigma-Aldrich)), and disrupted through sonication. Protein extracts were collected from the supernatant after centrifugation. Samples were resolved by SDS-PAGE and transferred to a nitrocellulose membrane using the iBlot Dry Blotting System (Invitrogen), according to manufacturer’s instructions. Transferred proteins were confirmed by Ponceau staining (0.25% Ponceau S in 40% methanol and 15% acetic acid). The membrane was blocked for at least 1 hour at room temperature with 5% dry milk prepared in PBT, and subsequently incubated overnight with the primary antibody (rabbit anti-GFP (i3S core facility, 1:1000)) diluted in blocking solution, at 4°C. The membrane was then washed three times for 10 minutes with PBT, and incubated for 1 hour with the secondary antibody conjugated to HRP (Jackson Immuno Research, 1:5000) diluted in blocking solution, at room temperature. After washing the membrane again three times for 10 minutes with PBT, blots were developed with Clarity Western ECL Substrate (Bio-Rad) and detected on X-ray films (Fuji Medical).

### Data processing and analysis

Image processing and quantifications were done with FIJI (Schindelin *et al*, 2012). For representative midsagittal images from egg chambers, we used a single optical section or maximum intensity projections of 2-5 planes. Surface or bottom images of egg chambers correspond to maximum intensity projections of the sections encompassing the epithelial region of interest. For live imaging processing of egg chambers, two FIJI features were always applied: the *StackReg* plugin (EPFL; Biomedical Imaging Group), to correct for the egg chamber movement, and the *Gaussian Blur 3D* filter, to remove background noise. For live imaging of pupal notum, the *Bleach Correction* plugin was applied to correct bleaching.

Statistical analysis and graphs were generated using GraphPad Prism 9 (GraphPad Software, La Jolla, CA, USA).

#### Quantification of multinucleation rate for the Anillin RNAi modifier screen

For the *in vivo* genetic modifier screen, we scored the amount of multinucleated follicle cells in surface z-projections of non-proliferative stage 10 egg chambers (Δz = 1µm). We selected stage 10 egg chambers to evaluate the multinucleation ratio for two main reasons. First, follicle cells are no longer undergoing mitotic division, which allows us to directly associate the number of nuclei with the presence of a multinucleated cell. Second, follicle cells are larger at these stages, which facilitates automated segmentation of nuclei and cell membranes. To segment epithelial cells, we developed two automated macros for FIJI. The first macro projects a 1 µm thick section from the acquired z-stack centered around the nucleus; since nucleus position along the z-axis is variable due to egg chamber curvature, the optimal z-planes to project are determined locally based on the closest nucleus. To extract the multinucleation ratio, the second macro selects a central region of interest (ROI) of the epithelium (to avoid quantification in the borders of egg chambers whose nuclei/membrane could be missed from the projection), segments the cells (marked with Myristoylated-GFP) within this ROI, segments nuclei in the acquired image (marked with DAPI) and then counts the number of nuclei within the previously segmented cells (see Fig S1 for further details). Manual validation of the segmentation was performed for each image to correct any errors prior to the quantification of the nuclei/cell ratio (multinucleation ratio). In the context of the modifier screen (Fig 1D-G), the multinucleation ratio for each egg chamber was compared to the mean of a control group of cytokinesis-sensitized (Anillin RNAi) egg chambers expressing UAS-mCherry. This control group was incubated during the same period of time of each of the three independent experiments performed for each RNAi line. Δmultinucleation ratio for each egg chamber is calculated as (nuclei/cell)_cell adhesion RNAi_ - (1/n) * Σ(X(nuclei/cell)i)*_control_*, where n represents the number of control egg chambers within each replicate.

#### Quantification of multinucleation frequency in Dys mutants

Due to genetic constraints, we used a different genetically encoded marker to visualize the epithelial cell cortex (ECad:GFP) than the one used in the *in vivo* genetic modifier screen. The number of mono and multinucleated cells was quantified in cross-sections in non-proliferative stage 10 egg chambers. Similar to the approach used for the screen, this quantification was restricted to the central area of the follicular epithelium.

#### Quantification of ring constriction rate

Analysis of cytokinetic ring constriction was performed in egg chambers expressing a tagged version of non-muscle myosin light chain, Sqh:mKate2 (Fig 4D, E and G), or non-muscle myosin heavy chain, Zip:GFP (Fig 4F and G). The diameter of the contractile ring along the apical-basal axis of the epithelium was manually measured using FIJI, from cytokinesis onset (t_0_) to complete constriction (t_f_). To depict the change in ring diameter through time (Fig 4D and F), these values were then normalized to the length of the ring at t_0_ and plotted as a function of time, using Prism (GraphPad Software). To measure constriction rate during the linear phase of constriction (∼ 80%-40% of the original diameter), we selected four consecutive timepoints that could fit to a linear regression (R^2^>0.95 was set as a threshold for the linear fit to assume constant constriction rate).

#### Quantification of cytokinesis failure in time-lapse movies

To understand when epithelial cells failed cytokinesis upon Anillin depletion and/or *Dys* loss-of-function, time-lapse movies were generated in egg chambers expressing Zip:GFP and labelled with a plasma membrane marker. We limited the analysis to small cells to prevent misleading results from defects accumulated from prior cytokinesis failure.

Moreover, we only included in the analysis cells that were imaged at least 20 minutes after ring constriction. Cells were scored in three categories: 1) not failing cytokinesis, 2) failing cytokinesis during ring constriction or 3) failing cytokinesis post-constriction.

## ACKNOWLEDGEMENTS

We thank X. Wang, M.-L Parmentier, the Bloomington *Drosophila* Stock Center and the Vienna *Drosophila* Resource center for fly stocks, and C. Sunkel for support with supervision. Work in the E.M. lab was funded by National Funds through FCT—Fundação para a Ciência e a Tecnologia, I.P., under the projects PTDC/BIA-CEL/1511/2021 and UIDB/04293/2020, E.M. is funded by “FCT Scientific Employment Stimulus - Individual Call” program (CEECIND/00622/2017). M.G. was supported by a PhD fellowship from FCT (SFRH/BD/130708/2017) and by IPATIMUP through CANCER_CHALLENGE2022, which also supported C.L. We also acknowledge the support of the i3S Scientific Platform ALM, member of the national infrastructure PPBI - Portuguese Platform of Bioimaging (PPBI-POCI-01-0145-FEDER-022122) and the Institut Curie PICT-IBiSA@BDD imaging facility (member of the French National Research Infrastructure France-BioImaging, ANR-10-INBS-04). Work in the V.M. lab was supported by the Association Française contre les Myopathies (AFM-Téléthon) (MyoNeurAlp2 network) and by the French government IDEX-ISITE initiative 16-IDEX-0001 (CAP 20-25). Work in the Y.B. lab is supported by the Institut Curie, the CNRS, the INSERM as well as ARC (SL220130607097), ANR (TiMecaDiv 20CE13000801), CANCERO-INCA (PLBIO2020/BELLAICHE) grants.

## AUTHOR CONTRIBUTIONS

E.M. and M.G. conceptualized the study and wrote the original manuscript draft. Data acquisition: M.G., C.L., C.R. and E.M. performed all experiments in the follicular epithelium. M.O. generated code for multinucleation analysis. H.A., G.R. and V.M. generated all isoform-specific *Dys* mutant lines. F.B. and Y.B contributed with experiments in the notum epithelium. Visualization: E.M and M.G. Data analysis and interpretation: M.G., C.L., F.B., Y.B, V.M. and E.M. Supervision: Y.B., V.M. and E.M. Funding acquisition: Y.B., V.M and E.M. Manuscript revision: M.G., C.L, M.O., F.B, Y.B, V.M. and E.M.

