## Supplemental data for "The Dystrophin-Dystroglycan complex ensures cytokinesis efficiency in *Drosophila* epithelia"

### Figure S1. Pipeline for the quantification of cytokinesis defects in the follicular epithelium of stage 10 egg chambers

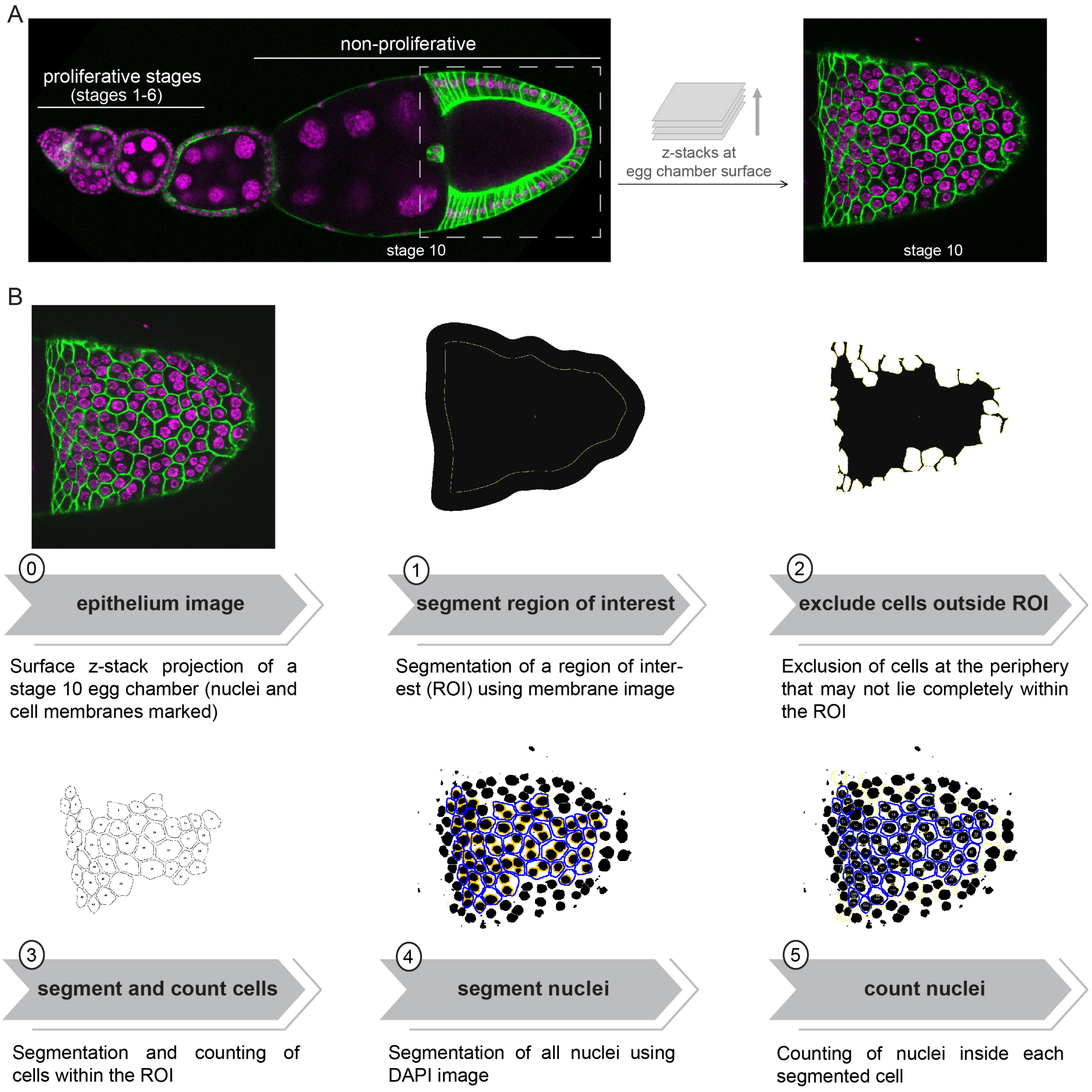

**A)** Surface z-projections of non-proliferative stage 10 egg chambers were used to quantify the cytokinesis defects from the *in vivo* genetic modifier screen (nuclei marked with DAPI (magenta) and cell membranes endogenously expressing Myr:GFP (green)). At this stage of oogenesis, follicle cells are no longer undergoing mitosis, allowing us to directly correlate multinucleation with cytokinesis failure, and are larger than at younger stages, facilitating the automated segmentation of nuclei and cell membranes. **B)** A Fiji macro was developed for the automated segmentation and counting of nuclei and cells in two-channel images of the follicular epithelium (nuclei in magenta and cell membranes in green). Note that this was restricted to the central area of the egg chamber, to avoid misleading quantifications from nuclei/cells at the periphery of the egg chambers. After running the macro, a manual validation of the segmentation of both nuclei and cells was performed for all images. The obtained values were used to calculate the Multinucleation Ratio, as explained in detail in the Materials and Methods section.

**Figure S2. Complementary analysis of RNAi-mediated depletion and their effects on tissue organization and cytokinesis (*related to Figure 1*)**

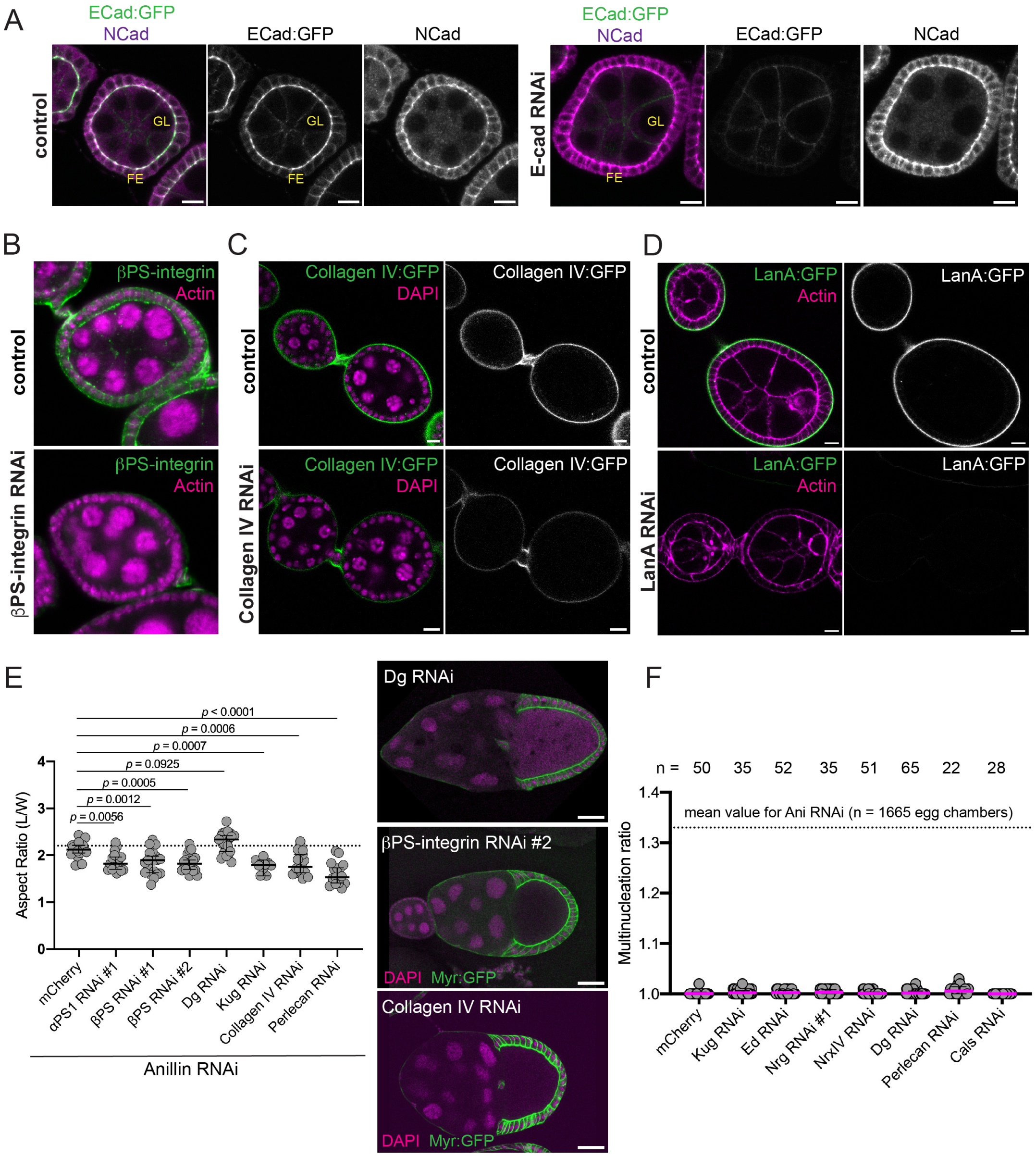

**A)** Midsagittal images of control and ECad RNAi egg chambers endogenously expressing ECad:GFP (green) and stained for NCad (magenta). Expression of RNAi for ECad in the follicular epithelium (FE) induces efficient protein depletion. As anticipated, reduction of ECad:GFP fluorescence by ECad RNAi is restricted to the follicular epithelium (FE) and ECad levels are normal in the germline (GL). Scale bar: 10 μm. **B)** *Midsagittal images of control and βPS-integrin* RNAi egg chambers, stained for *βPS-integrin (green) and F-actin (magenta). Expression of RNAi for βPS-integrin in the follicular epithelium induces protein depletion. Scale bar:* 10 μm. **C)** Midsagittal images of control and Collagen IV RNAi egg chambers endogenously expressing Collagen IV:GFP (green) and stained for DAPI (magenta). Collagen IV RNAi induces protein depletion in comparison with control. *Scale bar:* 10 μm. **D)** Midsagittal images of control and LanA RNAi egg chambers endogenously expressing LanA:GFP (green) and stained for DAPI (magenta). Expression of RNAi for LanA induces efficient protein depletion. *Scale bar:* 10 μm. **E)** Midsagittal images and measurement of the aspect ratio in stage 10 egg chambers co-depleted for Anillin and the indicated basal proteins, in comparison with control (UAS-mCherry). Aspect ratio was calculated as the ratio between egg chamber length and width, as depicted in the bottom image (DAPI in magenta and cell membrane in green (Myr:GFP)). Each dot represents the aspect ratio of an analyzed egg chamber. The dashed line represents the anticipated aspect ratio value for this egg chamber stage, as calculated in *Jia et al. 2016*. *p*-value was calculated by an ordinary one-way ANOVA. *Scale bar:* 50 μm. **F)** Multinucleated ratio in egg chambers depleted for the regulators of cell-cell and cell-matrix interactions that enhanced multinucleation in the Anillin RNAi modifier screen (Fig 1D-F), in comparison with control (UAS-mCherry). Depletion of these molecules on their own (in an unperturbed cytokinetic background) does not cause multinucleation. Each dot represents the Multinucleation Ratio of an analyzed egg chamber. Median is indicated. The dashed line represents the mean value of the multinucleation ratio for co-expression of Anillin RNAi with the control line (UAS-mCherry).

**Figure S3. Dynamic redistribution of the DAPC during epithelial cytokinesis in the *Drosophila* pupal notum**

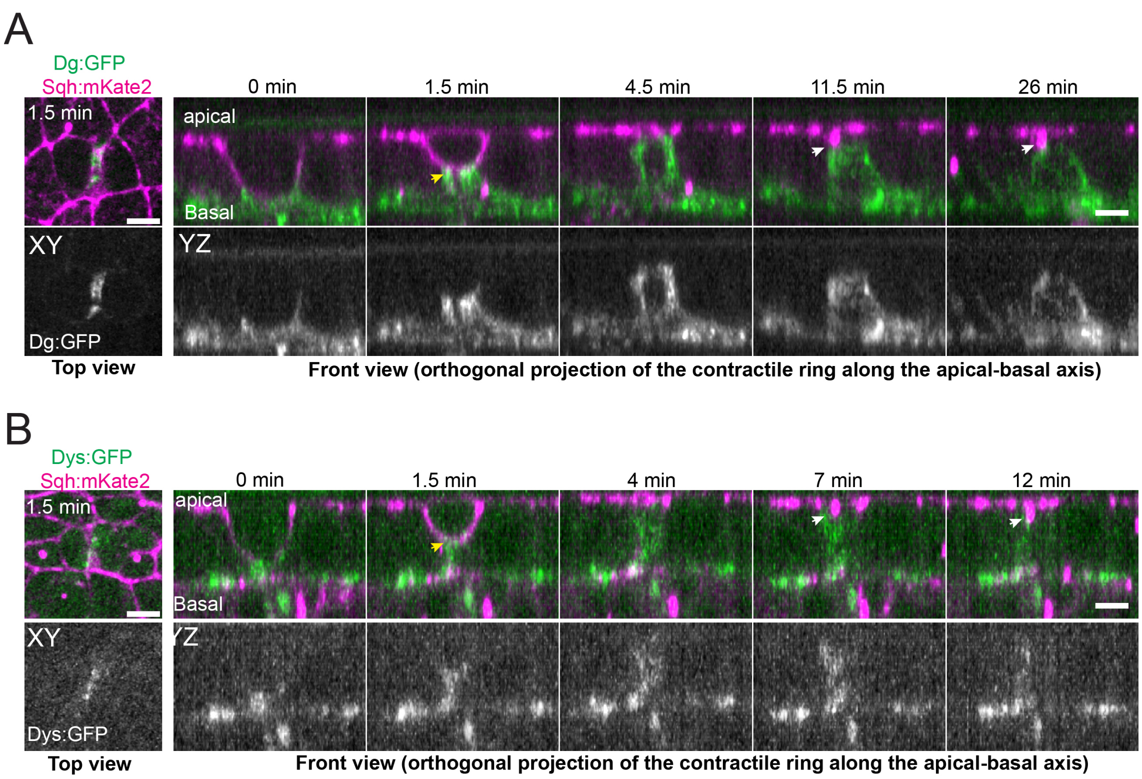

**A, B)** Redistribution of endogenously tagged Dg:GFP (A) and Dys:GFP (B) during epithelial cytokinesis in the *Drosophila* pupal dorsal thorax (notum). During ring (labelled with Sqh:mKate2) constriction, Dg:GFP (A) and Dys:GFP (B) accumulate at the basal part of the ingressing membrane (yellow arrows). After ring closure, both proteins become enriched close to the midbody (white arrows), similarly to what is observed in the *Drosophila* follicular epithelium. Scale bars: 5 µm.

### Figure S4. Spatial distribution of Dystrophin isoforms in stage 4 egg chambers

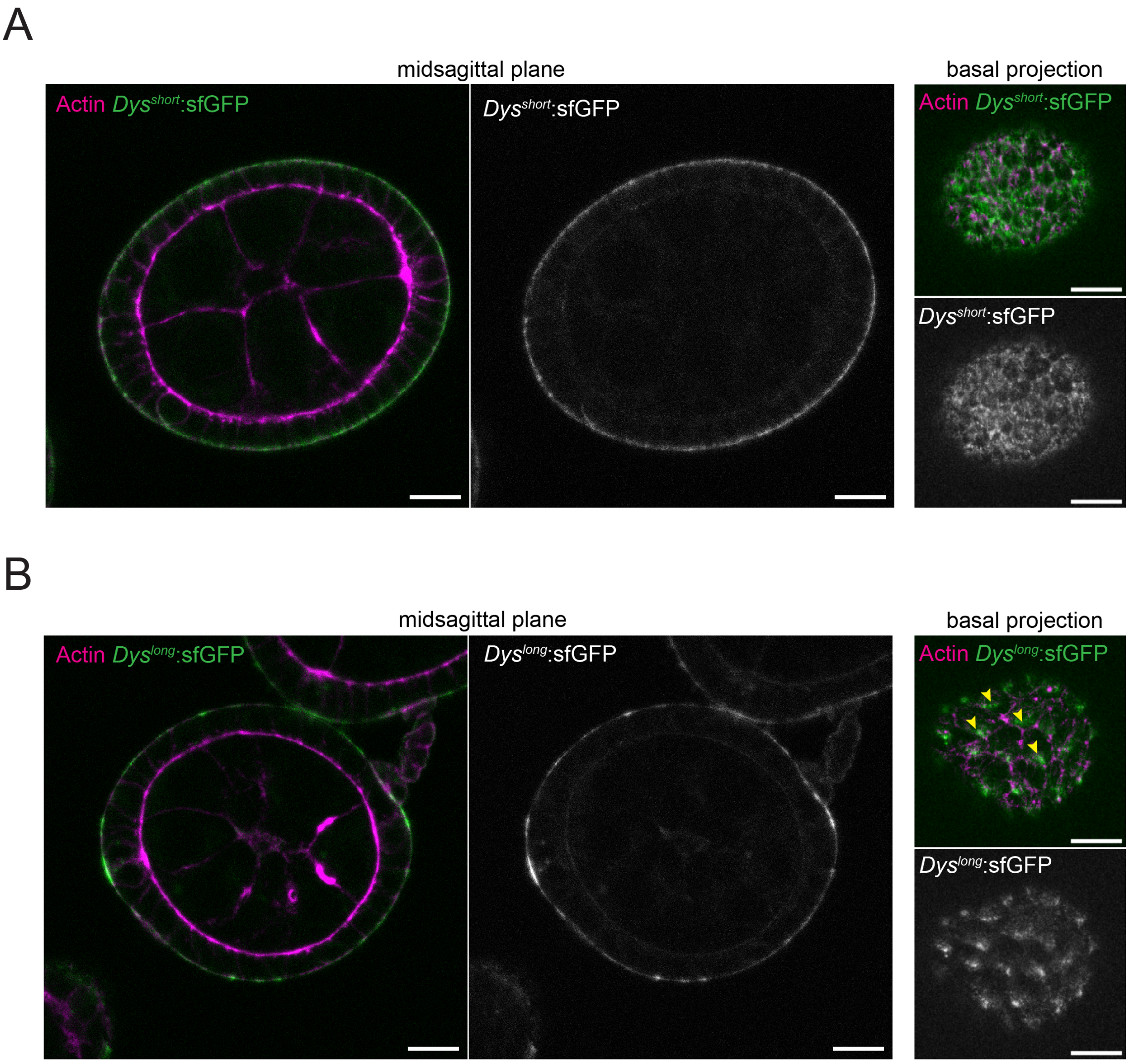

**A)** Representative midsagittal images (left) and basal projection (right) of the proliferative follicular epithelium endogenously expressing *Dys*$^{short}$:sfGFP (green) and stained for actin (magenta). *Dys*$^{short}$ mainly localizes at the basal domain of epithelial cells. Scale bar: 5 μm. **B)** Representative midsagittal images (left) and basal projection (right) of the proliferative follicular epithelium endogenously expressing *Dys*$^{long}$:sfGFP (green) and stained for actin (magenta). *Dys*$^{long}$ mainly localizes to the basal domain in a planar polarized manner. Scale bar: 5 μm.

### TABLES

#### Table S1 (xls). Screen overview

Table with an overview of the data from the *in vivo* modifier screen. The following information is provided (sub-divided by family of proteins): in column A, the name of each depleted protein and, in column B, the RNAi stock line used for protein depletion in the screen. In column C, the sample size (number of independent experiments, dissected egg chambers and quantified cells). In column D, the effect of protein depletion on the multinucleation ratio caused by co-depletion of Anillin. In column E, the median value and, in column F, the standard deviation of the multinucleation ratio (number of nuclei/number of cells) from all experiments for each depleted protein. In column G, the statistical significance of the *t*-test for each depleted protein against the respective experimental control (UAS-mCherry). In column H, the median value of the $\Delta$ multinucleation ratio (= $(\text{nuclei/cell})_{\text{RNAi}}$ - mean $(\text{nuclei/cell})_{\text{mCherry}}$) from all the replicates for each protein RNAi. In column I, the anticipated sub-cellular localization on the follicular epithelium of each protein and, in column J, the respective human orthologue(s).

**Table S2. Mutant alleles and transgenes**

| Mutant allele or transgene | Citation | Stock number or lab of origin |
|---|---|---|
| *Dys*$^{Df}$ | (Christoforou *et al*, 2008) | BDSC #7663 |
| *Dys*$^{E17}$ | (Christoforou *et al.*, 2008) | BDSC #63047 |
| *Dys*$^{long181}$ | this paper | Vincent Mirouse |
| *Dys*$^{RE225}$ | this paper | Vincent Mirouse |
| *Dg*$^{086}$ | (Christoforou *et al.*, 2008) | BDSC #63049 |
| *Dg*$^{043}$ | (Christoforou *et al.*, 2008) | BDSC #63048 |
| UAS-Dg:GFP | (Bogdanik *et al*, 2008) | Marie-Laure Parmentier |
| Dys:sfGFP | this study | Vincent Mirouse |
| *Dys*$^{short}$:sfGFP | this study | Vincent Mirouse |
| *Dys*$^{long}$:sfGFP | this study | Vincent Mirouse |
| Dys:GFP | (Nagarkar-Jaiswal *et al*, 2015) | BDSC #59782 |
| Dg:GFP | (Villedieu *et al.*, 2023) | Yohanns Bellaiche |
| GFP:aPKC | (Chen *et al*, 2018) | Daniel St Johnston |
| Zip:GFP | (Lowe *et al*, 2014) | DGGR #115082 |
| Ecad:GFP | (Pinheiro *et al.*, 2017) | Yohanns Bellaiche |
| Sqh:mKate2 | (Pinheiro *et al.*, 2017) | Yohanns Bellaiche |
| tub-Gal80$^{ts}$ | (McGuire *et al.*, 2003) | BDSC #7018 |
| *tj-Gal4* | (Olivieri *et al.*, 2010) | DGGR #104055 |
| UAS-Anillin RNAi | | VDRC #104674 |
| UAS-LARIAT | (Qin *et al.*, 2017) | Xiaobo Wang |
| UAS-Myr:GFP | | BDSC #32197 |
| *W*$^{1118}$ | | BDSC #3605 |

**Table S3. List of *Drosophila* genotypes**

| Figure | Genotype |
|---|---|
| 1B, 1C | *tj-Gal4*, UAS-Anillin RNAi/+; UAS-Myr:GFP, tub-Gal80$^{ts}$/UAS-mCherry<br>*tj-Gal4*, UAS-Anillin RNAi/UAS-ECad; UAS-Myr:GFP, tub-Gal80$^{ts}$/+ |
| 1D, 1E, 1F, 1G | *tj-Gal4*, UAS-Anillin RNAi/+; UAS-Myr:GFP, tub-Gal80$^{ts}$/UAS-mCherry<br>*tj-Gal4*, UAS-Anillin RNAi/CAM or ECM-protein RNAi; UAS-Myr:GFP, tub-Gal80$^{ts}$/+ <u>or</u> *tj-Gal4*, UAS-Anillin RNAi/+; UAS-Myr:GFP, tub-Gal80$^{ts}$/CAM or ECM-protein RNAi |
| 2B, 2C | *tj-Gal4*, ECad:GFP/+; *Dys*$^{E17/Df}$<br>*tj-Gal4*, ECad:GFP/UAS-Anillin RNAi<br>*tj-Gal4*, ECad:GFP/UAS-Anillin RNAi; *Dys*$^{E17/Df}$<br>*tj-Gal4*, ECad:GFP/+; *Dys*$^{MI025024/Df}$<br>*tj-Gal4*, ECad:GFP/UAS-Anillin RNAi; *Dys*$^{MI025024/Df}$ |
| 2E, 2F, 2G | *tj-Gal4*, UAS-Anillin RNAi/UAS-LARIAT; Dys:sfGFP |
| 3A | *tj-Gal4*, UAS-Dg:GFP/UASp-mRFP:Anillin |
| 3B | Dg:GFP/Sqh:mKate2 |
| 3C, 3D | Sqh:mKate2/+; Dys:sfGFP |
| 3E | GFP:aPKC/CyO; MKRS/TM6<br>IF/CyO; Dys:sfGFP<br>*Dys*$^{short}$:sfGFP(/TM3)<br>*Dys*$^{long}$:sfGFP(/TM3) |
| 3F | Sqh:mKate2/+; *Dys*$^{short}$:sfGFP<br>Sqh:mKate2/+; *Dys*$^{long}$:sfGFP |
| 3G, 3H | *tj-Gal4*, ECad:GFP/+; MKRS/TM6<br>*tj-Gal4*, ECad:GFP/+; *Dys*$^{long181/Df}$<br>*tj-Gal4*, ECad:GFP/UAS-Anillin RNAi<br>*tj-Gal4*, ECad:GFP/UAS-Anillin RNAi; *Dys*$^{long181/Df}$<br>*tj-Gal4*, ECad:GFP/+; *Dys*$^{RE225/Df}$<br>*tj-Gal4*, ECad:GFP/UAS-Anillin RNAi; *Dys*$^{RE225/Df}$ |
| 4A | *tj-Gal4*, Zip:GFP/UAS-Anillin RNAi<br>*tj-Gal4*, Zip:GFP/UAS-Anillin RNAi; *Dys*$^{E17/Df}$ |
| 4B | *tj-Gal4*, Zip:GFP/+<br>*tj-Gal4*, Zip:GFP/UAS-Anillin RNAi<br>*tj-Gal4*, Zip:GFP/UAS-Anillin RNAi; *Dys*$^{E17/Df}$ |
| 4C | *tj-Gal4*, Zip:GFP/UAS-Anillin RNAi<br>*tj-Gal4*, Zip:GFP/UAS-Anillin RNAi; *Dys*$^{E17/Df}$<br>*tj-Gal4*, Zip:GFP/UAS-Anillin RNAi; *Dys*$^{long181/Df}$ |

| | |
|---|---|
| | *tj-Gal4*, Zip:GFP/UAS-Anillin RNAi; *dys*$^{RE225/Df}$ |
| 4D, 4E | *tj-Gal4*, Sqh:mKate2/+<br>*Dg*$^{086/043}$; Sqh:mKate2/+<br>Sqh:mKate2/+; *Dys*$^{E17/Df}$ |
| 4F | *tj-Gal4*, Zip:GFP/+<br>*tj-Gal4*, Zip:GFP/+; *Dys*$^{long181/Df}$<br>*tj-Gal4*, Zip:GFP/+; *Dys*$^{R225/Df}$ |
| 4G | *tj-Gal4*, Sqh:mKate2/+<br>*Dg*$^{086/043}$; Sqh:mKate2/+<br>Sqh:mKate2/+; *Dys*$^{E17/Df}$<br>*tj-Gal4*, Zip:GFP/+<br>*tj-Gal4*, Zip:GFP/+; *Dys*$^{long181/Df}$<br>*tj-Gal4*, Zip:GFP/+; *Dys*$^{RE225/Df}$ |
| S1A, S1B | *tj-Gal4*, UAS-Anillin RNAi/+; UAS-Myr:GFP, tub-Gal80$^{ts}$/UAS-mCherry |
| S2A | *tj-Gal4*, ECad:GFP/+; UAS-mCherry RNAi/+<br>*tj-Gal4*, ECad:GFP/+; UAS-ECad RNAi/+ |
| S2B | *tj-Gal4*/+; UAS-mCherry/+<br>*tj-Gal4*/+; UAS-*βPS-integrin* RNAi/+ |
| S2C | *tj-Gal4*/Collagen IV:GFP; tub-Gal80$^{ts}$/+<br>*tj-Gal4*/Collagen IV:GFP; tub-Gal80$^{ts}$/UAS-Collagen IV RNAi |
| S2D | *tj-Gal4*/+; LanA:GFP/UAS-mCherry<br>*tj-Gal4*/+; LanA:GFP/UAS-LanA RNAi |
| S2E | *tj-Gal4*, UAS-Anillin RNAi/+; UAS-Myr:GFP tub-Gal80$^{ts}$/UAS-mCherry<br>*tj-Gal4*, UAS-Anillin RNAi/+; UAS-Myr:GFP tub-Gal80$^{ts}$/αPS1 RNAi #1<br>*tj-Gal4*, UAS-Anillin RNAi/+; UAS-Myr:GFP, tub-Gal80$^{ts}$/βPS RNAi #1<br>*tj-Gal4*, UAS-Anillin RNAi/+; UAS-Myr:GFP, tub-Gal80$^{ts}$/βPS RNAi #2<br>*tj-Gal4*, UAS-Anillin RNAi/+; UAS-Myr:GFP, tub-Gal80$^{ts}$/UAS-Dg RNAi<br>*tj-Gal4*, UAS-Anillin RNAi/UAS-Kug RNAi; UAS-Myr:GFP, tub-Gal80$^{ts}$/+<br>*tj-Gal4*, UAS-Anillin RNAi/+; UAS-Myr:GFP, tub-Gal80$^{ts}$/UAS-Collagen IV RNAi<br>*tj-Gal4*, UAS-Anillin RNAi/+; UAS-Myr:GFP, tub-Gal80$^{ts}$/UAS-Perlecan RNAi |
| S2F | *tj-Gal4*/+; UAS-Myr:GFP, tub-Gal80$^{ts}$/UAS-mCherry<br>*tj-Gal4*/UAS-Kug RNAi; UAS-Myr:GFP, tub-Gal80$^{ts}$/+<br>*tj-Gal4*/UAS-Ed RNAi; UAS-Myr:GFP, tub-Gal80$^{ts}$/+<br>*tj-Gal4*/+; UAS-Myr:GFP, tub-Gal80$^{ts}$/UAS-Nrg RNAi #1<br>*tj-Gal4*/+; UAS-Myr:GFP, tub-Gal80$^{ts}$/UAS-NrxIV RNAi<br>*tj-Gal4*/+; UAS-Myr:GFP, tub-Gal80$^{ts}$/UAS-Dg RNAi<br>*tj-Gal4*/+; UAS-Myr:GFP, tub-Gal80$^{ts}$/UAS-Perlecan RNAi<br>*tj-Gal4*/+; UAS-Myr:GFP, tub-Gal80$^{ts}$/UAS-Cals RNAi |
| S3A | Sqh:mKate2/Dg:GFP |
| S3B | Sqh:mKate2/+; Dys:GFP/+ |
| S4A | *Dys*$^{short}$:sfGFP/+ |
| S4B | *Dys*$^{long}$:sfGFP/+ |

**Table S4. Reagents**

| Reagent | Vendor (Cat#) |
|---|---|
| rat anti-N-cadherin antibody | DSHB (DN-Ex #8) |
| mouse anti-βPS-integrin antibody | DSHB (CF.6G11-s) |
| mouse anti-Armadillo antibody | DSHB (N2.7A1) |
| goat anti-rat Alexa 568 | Invitrogen (A11077) |
| goat anti-mouse Alexa 488 | Invitrogen (A11029) |
| goat anti-mouse Alexa 568 | Invitrogen (A11031) |
| Phalloidin-TRITC | Merck (P1951) |
| Vectashield Mounting Medium with DAPI | Vector Laboratories (H-1200-10) |
| CellMask Orange Plasma Membrane Stain | Thermo Fisher (C10045) |

**MOVIE LEGENDS**

**Movie S1. Dystrophin redistribution during epithelial cytokinesis.**
Time-lapse movie of an egg chamber expressing endogenous Dys:sfGFP (green) and Cell Mask Plasma Membrane stain (magenta) (left). An inset of a follicle cell dividing perpendicularly to the plane of imaging allows the visualization of Dys accumulating in the ingressing membrane during ring (labelled with Sqh:mKate2) constriction (right). Scale bar: 5 $\mu$m.

**Movie S2. Dystroglycan redistribution during epithelial cytokinesis.**
Time-lapse movie of an egg chamber expressing endogenous Dg:GFP (green) and Sqh:mKate2 (magenta) (left). An inset of a follicle cell dividing parallelly to the plane of imaging allows the visualization of Dys accumulating in the ingressing membrane, below the contractile ring (labelled with Sqh:mKate2) (right). Scale bar: 5 $\mu$m.

**Movie S3. Accumulation of Dystrophin at the newly-formed daughter cell interface.**
Time-lapse movie of surface projections of an egg chamber expressing endogenous Dys:sfGFP (green) and Sqh:mKate2 (magenta). After ring closure, Dys:sfGFP (middle) becomes enriched at the new daughter cell interface. Scale bar: 5 $\mu$m.

**Movie S4. Imaging of Dystrophin isoforms during epithelial cytokinesis.**
Time-lapse movies of dividing follicle cells endogenously expressing $Dys^{short}$:sfGFP (top) or $Dys^{long}$:sfGFP (bottom) (green) and Sqh:mKate2 (magenta). The dividing plane (parallel to the plane of imaging) allows for an end-on view of the contractile ring (labelled with Sqh:mKate2). Both isoforms display a local enrichment at the ingressing membrane during ring constriction, and close to the midbody (at the apical side of follicle cells) after ring closure. Scale bar: 5 $\mu$m.

**Movie S5. Dys contributes for the efficiency of the last stage of cytokinesis.**
Time-lapse movie of surface projections of dividing follicle cells endogenously expressing Zip:GFP (green) and stained for a membrane marker (magenta), in *Dys* mutant egg chambers depleted of Anillin by RNAi. Cells fail cytokinesis due to membrane regression after ring closure. Scale bar: 5 $\mu$m.

**Movie S6. DAPC ensures normal contractile ring constriction in the follicular epithelium.**
Time-lapse movies of cytokinetic ring constriction viewed along the apical-basal axis in control, $Dg^{086/043}$ and $Dys^{Df/E17}$ mutant egg chambers, expressing Sqh:mKate2. Disruption of either Dg or Dys function delays ring constriction in the follicular epithelium. Scale bar: 5 $\mu$m.
